# TDP-43 regulates GAD1 mRNA splicing and GABA signaling in Drosophila brains

**DOI:** 10.1101/2021.04.16.440162

**Authors:** Giulia Romano, Nikola Holodkov, Raffaella Klima, Fabian Feiguin

## Abstract

Alterations in the function of the RNA-binding protein TDP-43 is largely associated with the pathogenesis of amyotrophic lateral sclerosis (ALS), a devastating disease of the human motor system that leads to motoneurons degeneration and reduced life expectancy by molecular mechanisms not well known. Regarding to that, we found that the expression levels of the glutamic acid decarboxylase enzyme (GAD1), responsible to convert glutamate to γ-aminobutyric acid (GABA), were downregulated in TBPH-null flies and motoneurons derived from ALS patients carrying mutations in TDP-43 suggesting that defects in the regulation of GAD1 may lead to neurodegeneration by affecting neurotransmitter balance. In this study, we observed that TBPH was required to regulate GAD1 pre-mRNA splicing and GABA levels in Drosophila brains. Interestingly, we discovered that pharmacological treatments aimed to modulate GABA neurotransmission were able to revert locomotion deficiencies in TBPH-minus flies revealing novel mechanisms and therapeutic strategies in ALS.

## INTRODUCTION

A common characteristic shared by several neurodegenerative diseases is the dysfunction of the RNA-binding protein TDP-43, a member of the heterogenous nuclear ribonucleoproteins (hnRNPs) family (Cook et al., 2008; de Boer et al., 2021). Initially identified as the main component of the ubiquitinated cytoplasmic inclusions in ALS and frontotemporal lobar degeneration (FTLD) (Arai et al., 2006; Geser et al., 2009; Neumann et al., 2006), TDP-43 pathology is currently described in a large proportion of cases of Alzheimer’s disease as well as Parkinson’s and Huntington’s disease (Amador-Ortiz et al., 2007; Higashi et al., 2007; Josephs et al., 2014; Schwab et al., 2008). Attentive studies aimed to understand the normal function of TDP-43 and its participation in the mechanisms of neurodegeneration have, therefore, became critical to establish the metabolic pathways implicated in TDP-43-mediated neuronal toxicity. In this direction we have previously indicated that TDP-43 is required to regulate the synaptic levels of GAD1, the enzyme responsible to convert glutamate to γ-aminobutyric acid (GABA), suggesting that modifications in the glutamate/GABA neurotransmitter balance may affect neuronal survival in TDP-43 perturbed brains (Romano et al., 2018). If the involvement of glutamate in the pathogenesis of ALS disease was already suggested by several studies (Foran and Trotti, 2009; Trotti et al., 2001; Van Den Bosch et al., 2006), GABA implications are less known. Consequently, in this study we investigated the mechanisms by which TDP-43 regulates the cytoplasmic levels of GAD1 and determined the role of GABA neurotransmission in TDP-43 disease using Drosophila.

## MATERIALS and METHODS

### Fly strains

The fly genotypes used for the experiments are indicated hereafter: w^1118^ – OregonR – w;tbph^Δ23^/CyO^GFP^ – w;tbph^Δ142^/CyO^GFP^ – w;*elav*-GAL4/CyO^GFP^ – w,UAS-Dicer2;*elav*-GAL4/CyO^GFP^ – w;siRNA-UAS-GFP (#9330, BDSC) – w;UAS-GAD1/TM3,Sb (gifted from Dr. Andreas Prokop) – w;UAS-TBPH — w;UAS-TBPH-RNAi/TM6B (#38377, VDRC) – siRNA-GABA_A_ receptor, resistant to dieldrin (RDL, #41101, #100429, VDRC) – siRNA Ligand Gated Chloride Channel (LCCH3, #37409, #109606, VDRC) – siRNA Glycine like receptor (GRD, #38384, #58175) – siRNA Vesicular GABA Transporter (vGAT, #45916, VDRC) – siRNA-GABA_B_ Receptor Type 1 (R1, #101440, VDRC) – siRNA-GABA_B_ receptor Type 2 (R2, #1784, #1785, #110268, VDRC) – siRNA-GABA_B_ receptor Type 3 (R3, #50622, #108036, VDRC).

### Climbing assay

Freshly eclosed flies were transferred in new food vials and let to adapt for 3 to 4 days. After this period, they were moved, without anesthesia, to 15 ml glass cylinders, tapped to the bottom and let climb taking advantage of their natural drive for negative geotaxis. The number of flies that reached the top of the tube in 15 s was counted, and converted in percentage.

### Larval Brain, Microscope Acquisition and Quantification

Larvae were selected and dissected as previously described in (Romano et al., 2014). Briefly, for whole larval brain staining, previously selected larvae were dissected on Sylgard plates in Phosphate Buffer (PB), brains were removed and fixed 20 min in 4% formaldehyde in Phosphate Buffer with 0.3% Triton X100 (PBT), blocked in 5% Normal Goat Serum (NGS), labelled with primary and secondary antibodies and mounted on microscope slides in Slow Fade (S36936, Thermofisher). Brain images were acquired on 20x at a 0.6-fold magnification on a LSM 880 Zeiss confocal microscope and then analyzed using ImageJ. Z-stacking was performed and the ratio of GABA/Elav Max Intensity was calculated. Primary antibodies concentration: anti-Elav (1:250, DSHB), anti-GABA (1:500, Sigma). Secondary antibodies concentration: Alexa Fluor 488 (1:500, Life Technologies), Alexa Fluor 555 (1:500, Life Technologies).

### Cell culture and RNA interference

S2 cells were cultured in Insect-Xpress medium (Lonza) supplemented 10% fetal bovine serum and 1X antibiotic-antimycotic solution (#A5955, Sigma). RNA interference was achieved using HiPerfect Transfection Reagent (#301705, Qiagen) and siRNA specific for Drosophila TBPH (5’-GGAAGACCACAGAGGAGAGC-3’), as control siRNA for luciferase was used (5’-UAAGGCUAUGAAGAGAUAC-3’; Sigma). Immediately before transfection 2×10^6^ cells were seeded in 6-well plates in 1.4 ml of medium containing 10% fetal serum. 2μg of each siRNA, were added to 91μl of Opti-MEM I reduced serum medium (#51985-026, Thermo Fisher Scientific), incubated 5 minutes at room temperature and subsequently 6μl of HiPerfect Transfection Reagent were added. The silencing procedure was performed again after 24 hours. Plasmid containing the Gad1 mini-gene and Gal4 were co-transfected (0,3μg each) at 24 hours together with the second siRNA dose. Cells were analyzed after 72 hours of the initial treatment.

### GAD1 minigene construction

A mini-gene containing the 5’ UTR region of Gad1 spanning from exon1 to exon2 (ref isoform FBtr0073276) in frame with a reporter EGFP has been cloned in pUAST attB plasmid. The minigene was co-transfected with pGAL4 in S2 cells. The EGFP expression was analyzed in S2 cells treated with siRNAs against TDP-43 or luciferase.

### RT-PCR

Adult heads were mechanically squeezed to proceed with RNA extraction, using RNeasy^®^Microarray tissue kit (#73304 Qiagen) and treated with Turbo DNA-free kit (#AM1907 Ambion). Retro-transcription was performed using oligo-dT primers and SuperScript^®^ III First-Strand Synthesis (#18080093 Invitrogen). Gene specific primers were designed for amplification:

Gad1(intronic set): *5’GCCAAACTCCGCATTCCATTT3’ and 5’ACCGAAGGCCGTGGGTCCTCG3’*

Gad1(exon/exon set): *5’TGATCCTTGAACCGGAGTGC3’ and 5’ACCATCAGCGTTCCCTTCTG3’*

Rpl11 *5’CCATCGGTATCTATGGTCTGGA3’ and 5’CATCGTATTTCTGCTGGAACCA3’*

The quantification was calculated according the ΔΔC_T_ equation and then normalized on control genotype.

### Western Blot

S2 cells were lysed in lysis buffer (10mM Tris, 150 mM NaCl, 5 mM EDTA, 5 mM EGTA, 10% Glycerol, 50 mM NaF, 5 mM DTT, 4 M Urea, pH 7.4, protease inhibitors (Roche, #11836170001)) and after protein quantification of lysates by Qbit (Q#33211, Invitrogen), samples were separated on 10% SDS-PAGE and transfer to 0,22μm nitrocellulose membranes (#NBA083C Whatman Protran). Membranes were blocked in 5% non-fat dry milk in Tris Buffered Saline with 0.1 % Tween-20 (TBS-T). Primary antibodies were used with the following concentrations: anti-GFP (1:2000, Invitrogen), anti-TBPH (1:3000, homemade (Feiguin et al., 2009) and anti - tubulin DM1A (1:5000, Calbiochem). Secondary antibodies concentration: anti-rabbit HRP conjugated (1:10000, Pierce #32460), anti-mouse HRP conjugated (1:10000, Pierce # 32430). For detection, SuperSignal™ West Femto Maximum Sensitivity Substrate Kit (Pierce, #PR34095) was used. Quantification was performed by ImageJ after acquisition of the autoradiographic films with an Epson Expression 1680 Pro Scanner.

### Drug treatment

Muscimol 100 μM and Nipecotic Acid 200 μM were used. According to the final concentration needed, drugs were prepared and diluted in the fresh fly food. This solution was aliquoted fly tubes where parental flies were transferred and let lay embryos for a period of 24 h. After this period, parental flies were discarded, and embryos were let to grow to third instar larvae, used for further analysis.

### Larval movement

Third instar larvae were individually selected, cleaned with water from any food remaining and placed on 0.7% agarose plates under stereoscope vision. After 30 s of adaptation, the number of peristaltic waves performed in 2 min were counted.

### Statistical Analysis

All statistical analysis was performed with GraphPad Prism 7.0 applying one-way ANOVA test (Bonferroni post-test).

## RESULTS

### Drosophila TDP-43 regulates the expression levels of GAD1 by facilitating its pre-mRNA splicing

We have previously demonstrated that the protein levels of GAD1 appeared downregulated in TBPH-null flies, however, the mechanisms behind these modifications were not identified (Romano et al., 2018). In order to address this question, and considering that TBPH is involved in the regulation of different aspects of the RNA metabolism, we decided to investigate whether the GAD1 RNA splicing was affected in TBPH-minus alleles. To this purpose, we designed a set of primers specific to discriminate GAD1 pre m-RNA and another one specific for m-RNA (respectively the intronic set and the exonic set, (see Fig. 1B), and quantified their expression levels by qRT-PCR. Interestingly, we found that the pre-mRNA of GAD1 appeared upregulated in the brains of the TBPH-mutant alleles compared to controls (Fig. 1B, left graph). On the contrary, the mature mRNA transcript of GAD1 was downregulated in the mutant flies judged against controls (Fig. 1B, right graph) indicating that TBPH is required to regulate the processing of GAD1 mRNA, most probably, through the splicing of the mature transcript in Drosophila brains. In agreement with this possibility, we observed the presence of putative binding sites for TBPH in the long intron present at the 5’UTR region of GAD1, between the non-coding exon1 and the coding exon2 (Fig.1C, upper scheme). In order to test if the absence of TBPH influenced the processing of these segments, we constructed a minigene containing the genomic sequences described above (Fig.1C, upper left scheme). The GAD1 pre-mRNA 5’-UTR minigene also presented an EGFP cloned in frame with the second exon of GAD1 that carried the original ATG starting codon. The construct was placed under the control of the GAL4-UAS system and used to transfect Drosophila S2 cells under the Actin promoter (actin-GAL4). Thus, S2 cells co-transfected with an RNAi against TBPH showed a strong reduction in the expression of the EGFP protein compared to control cells treated with an RNAi against luciferase in western blot assays (Fig.1C right panel, quantified in the left graph). These results show that TBPH function is required for the proper splicing and expression of the GAD1 pre-mRNA 5’-UTR minigene and strongly suggest that similar alterations may explain the defects detected in the processing of the immature GAD1 mRNA in TBPH-null flies.

**Figure 1.**
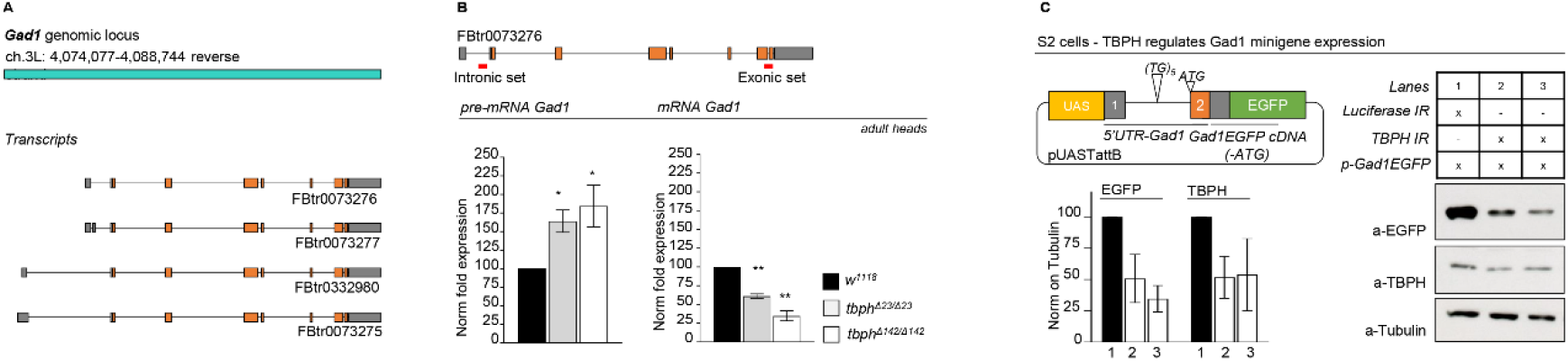
TBPH modulates GAD1 protein expression by regulating its pre-mRNA 5’-UTR splicing. **A** Scheme of Gad1 genomic locus and transcripts in *Drosophila*. **B** qRT-PCR of Gad1 pre-mRNA left panel and mRNA right panel on adult heads of w^1118^, tbph^Δ23/Δ23^ and tbph^Δ142/Δ142^. *n*=3; *p<0.05 and ** p<0.01 calculated by one-way Anova. **C** Western blot analysis for GAD1 minigene expression in S2 cells (anti-EGFP, anti-TBPH, and anti-Tubulin). Lane 1: S2 cells co-transfected with GAD1 minigene, Actin-GAL4 and Luciferase-IR. Lanes 2 and 3: S2 cells co-transfected with GAD1 minigene, Actin-GAL4, and TBPH-IR. In the graph the quantification of EGFP and TBPH expression weighted on Tubulin of Luciferase-IR (black column) compared with TBPH-IR (white columns). In the scheme GAD1 minigene: the 5’ UTR region of Gad1 spanning from exon1 to exon2 in frame with a reporter EGFP has been cloned in pUAST attB plasmid. *n*=2, Error bars SEM.

### Reduced levels of GABA neurotransmitter in TDP-43-null brains affect locomotive behaviors

The alterations in the processing of GAD1 immature mRNA described above insinuate that GABA neurotransmission might be affected in TBPH-null flies. In fact, the synthesis of GABA occurs through the conversion of glutamate by GAD1 enzyme; then GABA is loaded into vesicles by the vesicular GABA transporter (vGAT) (Owens and Kriegstein, 2002). The released GABA in the synaptic cleft is bind by GABA receptors placed on the post synaptic cell. The GABA receptors are divided into two types: the ionotropic and the metabotropic. To date in *Drosophila* three ionotropic subunits and three metabotropic subunits have been described (see Table 1). In order to address this hypothesis, we decided to test whether modifications in GABA signaling were able to influence the locomotive phenotypes described in TBPH deficient flies. For these experiments, we took advantage of the GAL4-UAS expression system to simultaneously co-silence the endogenous TBPH protein together with the various GABA ionotropic and metabotropic receptors.

**Table 1.**
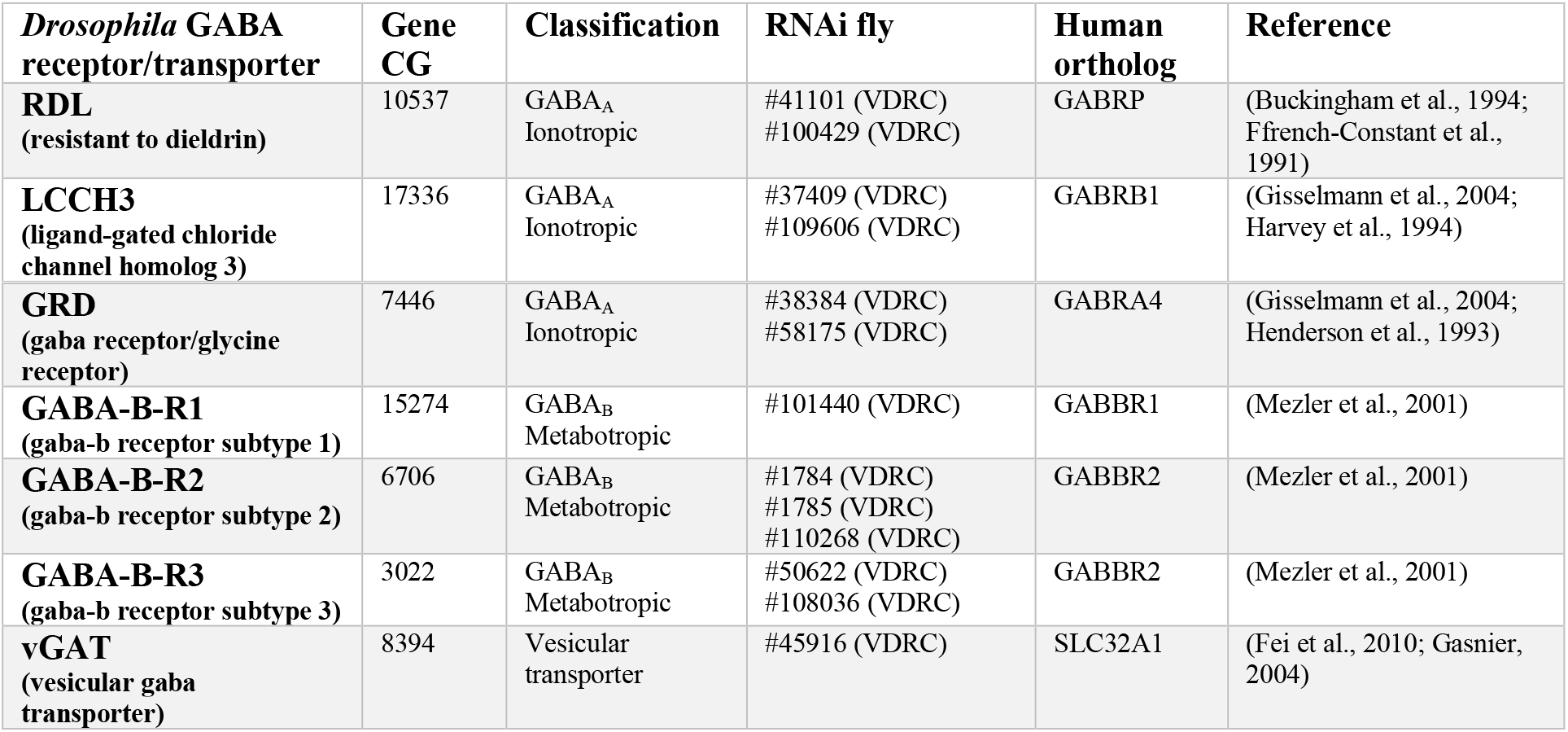
GABA receptors and transporters.

| *Drosophila* GABA receptor/transporter | Gene CG | Classification | RNAi fly | Human ortholog | Reference |
| --- | --- | --- | --- | --- | --- |
| **RDL**<br>(resistant to dieldrin) | 10537 | GABA<sub>A</sub><br>Ionotropic | #41101 (VDRC)<br>#100429 (VDRC) | GABRP | (Buckingham et al., 1994;<br>Ffrench-Constant et al., 1991) |
| **LCCH3**<br>(ligand-gated chloride channel homolog 3) | 17336 | GABA<sub>A</sub><br>Ionotropic | #37409 (VDRC)<br>#109606 (VDRC) | GABRB1 | (Gisselmann et al., 2004;<br>Harvey et al., 1994) |
| **GRD**<br>(gaba receptor/glycine receptor) | 7446 | GABA<sub>A</sub><br>Ionotropic | #38384 (VDRC)<br>#58175 (VDRC) | GABRA4 | (Gisselmann et al., 2004;<br>Henderson et al., 1993) |
| **GABA-B-R1**<br>(gaba-b receptor subtype 1) | 15274 | GABA<sub>B</sub><br>Metabotropic | #101440 (VDRC) | GABBR1 | (Mezler et al., 2001) |
| **GABA-B-R2**<br>(gaba-b receptor subtype 2) | 6706 | GABA<sub>B</sub><br>Metabotropic | #1784 (VDRC)<br>#1785 (VDRC)<br>#110268 (VDRC) | GABBR2 | (Mezler et al., 2001) |
| **GABA-B-R3**<br>(gaba-b receptor subtype 3) | 3022 | GABA<sub>B</sub><br>Metabotropic | #50622 (VDRC)<br>#108036 (VDRC) | GABBR2 | (Mezler et al., 2001) |
| **vGAT**<br>(vesicular gaba transporter) | 8394 | Vesicular<br>transporter | #45916 (VDRC) | SLC32A1 | (Fei et al., 2010; Gasnier,<br>2004) |

Thus, specific RNAi against the GABA_A_ type receptor (RDL), the ligand-gated chloride channel (LCCH3), the GABA_B_ type receptor 1, 2 and 3 (R1, R2 and R3) were expressed, either alone or together with an RNAi against TBPH, under the neuronal driver *elav*-GAL4. As a result, we found that the suppression of the GABA receptors strongly enhanced the motility problems present in TBPH-defective flies compared to control insects expressing the RNAi against the GABA receptors alone (Fig. 2 and Supplementary Figure 1).

**Figure 2.**
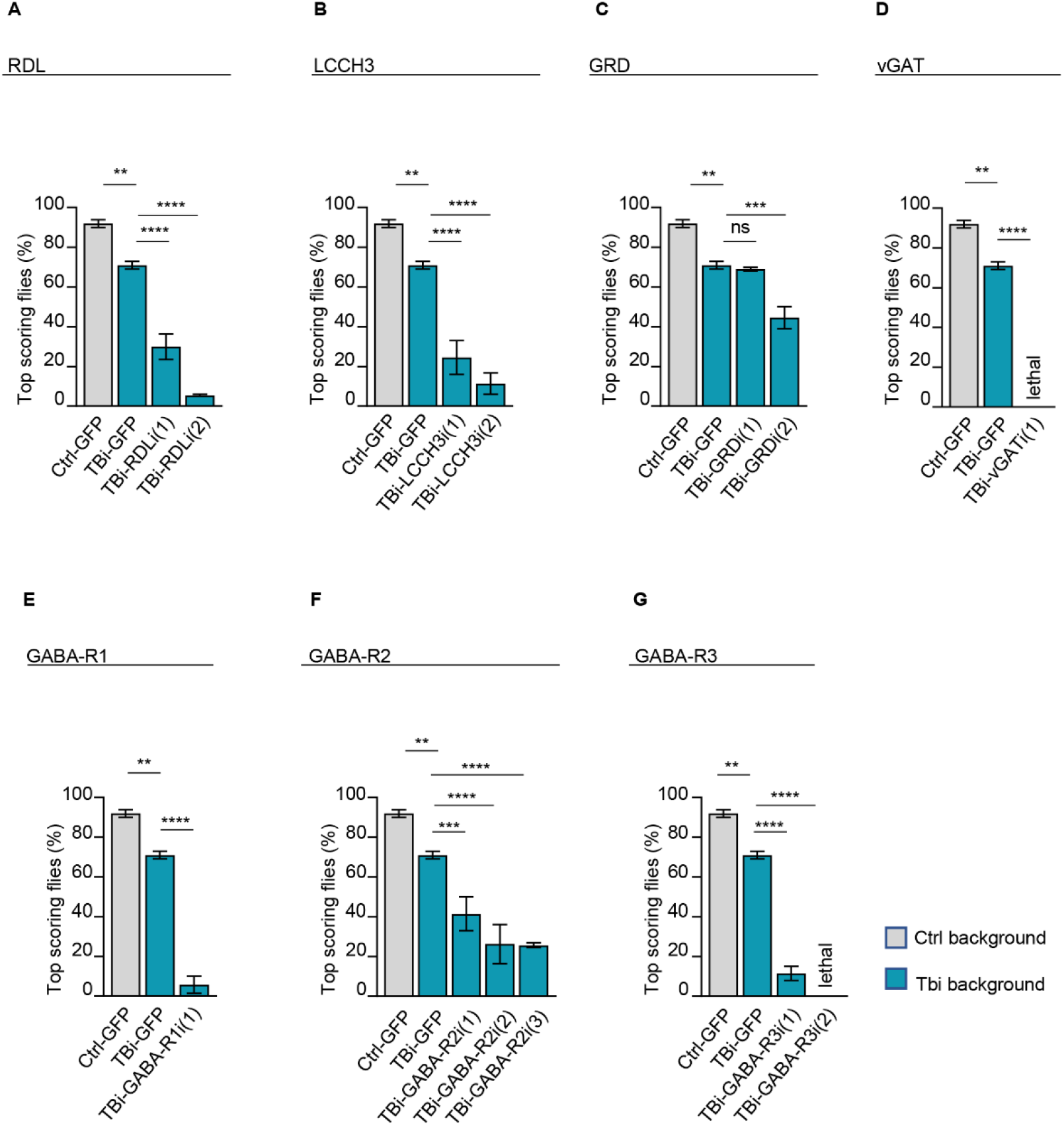
Silencing of GABA receptors and transporter worsen the TBPH hypomorphic phenotype. **A-G** Climbing assay of 4 days adult flies of control (Ctrl-GFP=tbph^Δ23^,*elav*-GAL4/UAS-GFP;UAS-Dicer/+), TBPH hypomorphic (TBi-GFP= tbph^Δ23^,*elav*-GAL4/UAS-GFP;UAS-Dicer/UAS-TBPH-IR) and TBPH hypomorphic with several GABA receptors silenced (TBi-GABA receptors name= tbph^Δ23^,*elav*-GAL4/UAS-GABA-receptor IR;UAS-Dicer/UAS-TBPH-IR). In **A** RDL-IR line (1)= #41101, line (2) #100429; in **B** LCCH3-IR line (1) #37409, line (2) #109606; in **C** GRD-IR line (1) #38384, line (2) #58175; in **D** vGAT-IR line (1) #45916; in **E** GABA_B_ receptor type 1 line (1) #101440; in **F** GABA_B_ receptor type 2 line (1) #1784, line (2) #1785, line (3) #110268; in **G** GABA_B_ receptor type 3 line (1) #50622, line (2) #108036. The total number of tested animals per genotype was > 50. ns=not significative, *p < 0.05, **p<0.01, ***p < 0.001 and ****p<0.0001; calculated by one-way ANOVA. Error bars SEM.

In the same direction, we utilized a specific antibody against GABA to quantify the intracellular levels of the neurotransmitter in third instar larval brain. After the staining, we found that the levels of GABA intensity appeared significantly reduced in TDP-43-null brains compared to wild type controls (Fig. 3A-B, E). Interestingly, we found that the neuronal transgenic expression of GAD1 or the endogenous protein TBPH in TDP-43-minus backgrounds were able to recover the expression levels of GABA in Drosophila brains (Fig. 3C-D, E), indicating that these results are specific and the regulation of GAD1 levels is critical to prevent GABA neurotransmission defects and neurodegeneration in TDP-43-defective brains.

**Figure 3.**
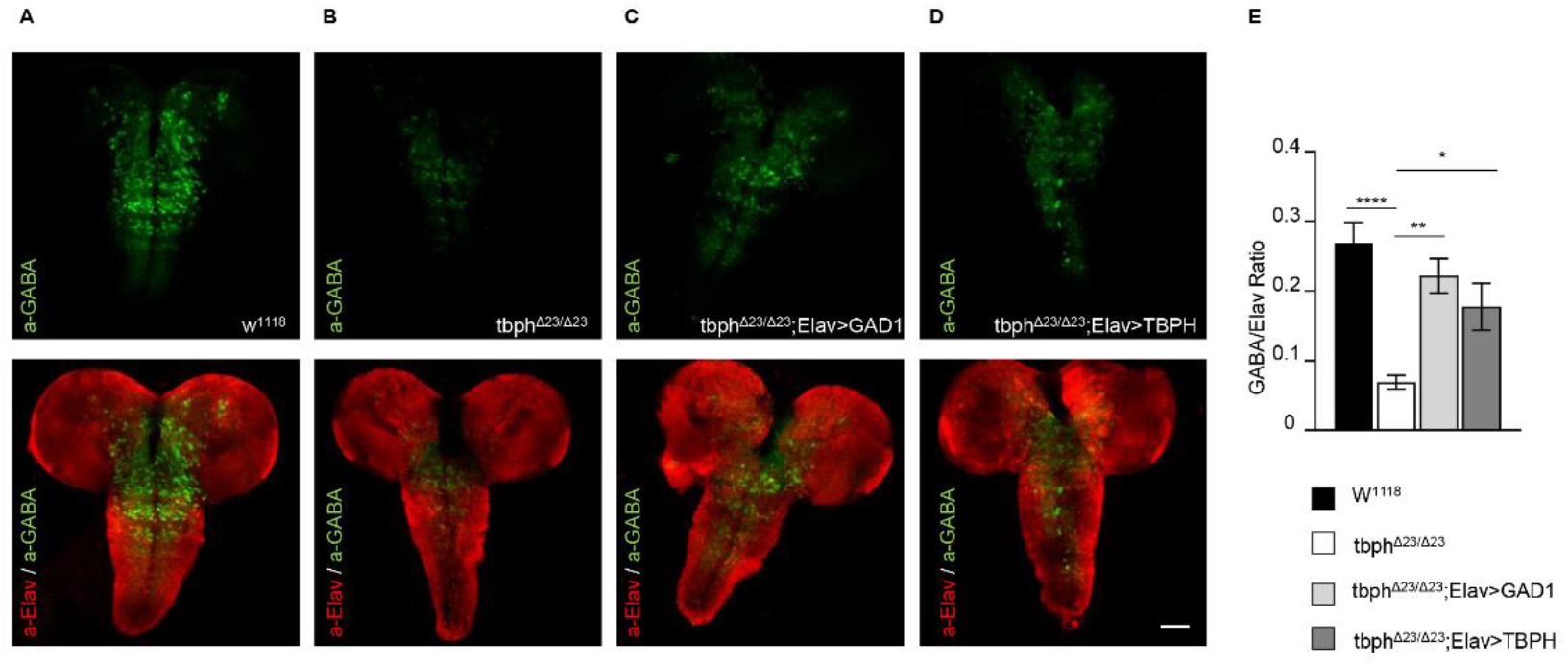
GABA was strongly downregulated in TBPH minus larval brains and recovered by GAD1 and TBPH expression in neurons. **A-D** Confocal microscopy acquisition of whole larval brains. GABA neurotransmitter was labeled with anti-GABA (Green) and neurons were labeled anti-Elav (Red, merged to GABA) of w^1118^ (**A**), tbph^Δ23/Δ23^ (**B**), tbph^Δ23^,*elav*-GAL4/tbph^Δ23^;UAS-GAD1/+ (**C**), and tbph^Δ23^,*elav*-GAL4/ tbph^Δ23^, UAS-TBPH (**D**). **E** Quantification of whole brain intensity GABA/Elav ratio. The number of examined brains per genotype was >10. *p < 0.05, **p < 0.01, ****p < 0.0001 calculated by one-way ANOVA. Error bars SEM. Scale bar: 50 μm.

### The recovery of GABA neurotransmission improves locomotion in TDP-43-null flies

In order to determine if the reduced levels of GABA detected in TBPH-minus brains were related with the neurodegenerative phenotypes described in these flies, we decided to treat TBPH-null flies with different agonists of GABA neurotransmission. For these assays, a potent GABA uptake inhibitor, Nipecotic acid (200 μM) was added to the fly food during larvae development and found that the administration of this compound was able to consistently improve the peristaltic movements of the TBPH-minus third-instar larvae (L3) compared to untreated mutants or wild-type controls (Fig. 4A).

**Figure 4.**
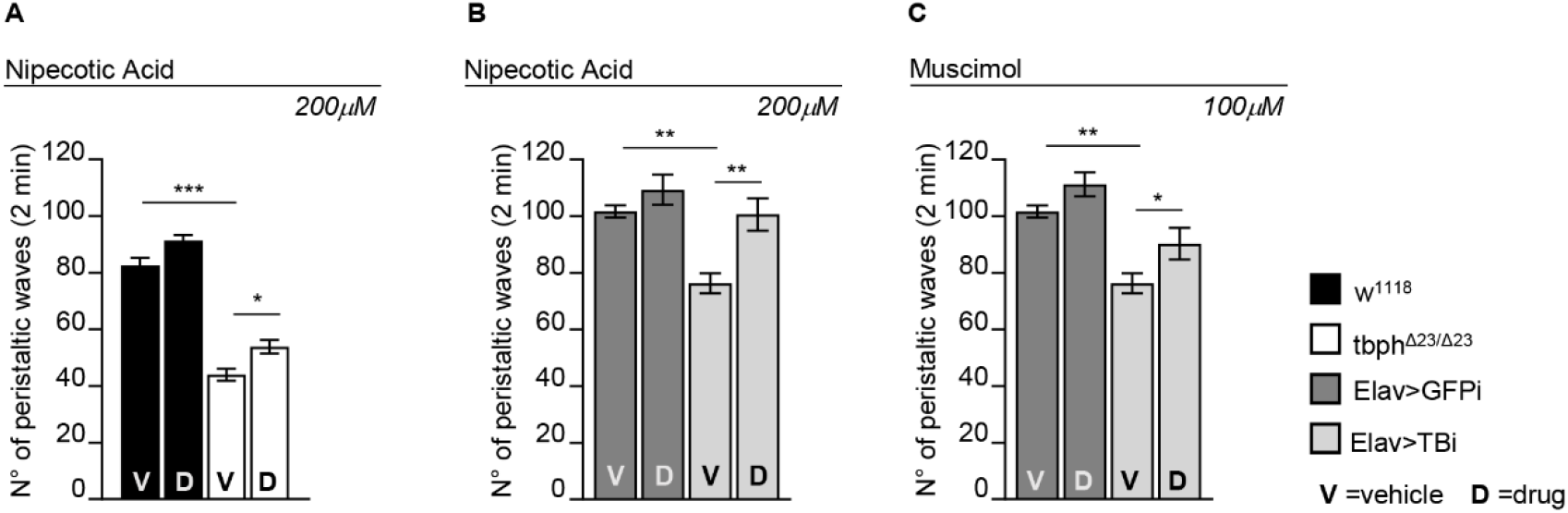
Pharmacological GABA stimulation ameliorates the motility defects in TBPH dysfunction. **A** Peristaltic larval waves of third instar larvae of w^1118^ (black columns) and tbph^Δ23/Δ23^ (white columns) fed with 200 μM of Nipecotic Acid (D) and vehicle only (V). **B** Peristaltic larval waves of third instar larvae of Elav>GFPi (w,UAS-Dicer/+; tbph^Δ23^,*elav*-GAL4/UAS-GFP-IR) (dark gray columns) and Elav>TBi (w,UAS-Dicer/+; tbph^Δ23^,*elav*-GAL4/+;UAS-TBPH-IR/+) (light gray columns) fed with 200 μM of Nipecotic Acid (D) and vehicle only (V). **C** Peristaltic larval waves of third instar larvae of Elav>GFPi (w,UAS-Dicer/+; tbph^Δ23^,*elav*-GAL4/UAS-GFP-IR) (dark gray columns) and Elav>TBi (w,UAS-Dicer/+; tbph^Δ23^,*elav*-GAL4/+;UAS-TBPH-IR/+) (light gray columns) fed with 100 μM of Muscimol (D) and vehicle only (V).n=20, *p < 0.05, **p<0.01, ***p<0.001, calculated by one-way ANOVA. Error bars SEM.

The positive effect of this pharmacological treatment became more obvious when TBPH hypomorphic background was utilized. As a matter of fact, we found that 200μM of Nipecotic Acid dispensed to flies expressing TBPH-RNAi in neurons (UAS-Dcr-2/+;tbph^Δ23^,*elav*-GAL4/+;TBPH-RNAi/+) were sufficient to recover motility (Fig. 4B), indicating that GABA neurotransmission plays an important role in TBPH-mediated neurodegeneration. Similar approach utilizing a different drug, the Muscimol (100 μM) (a known GABA_A_ receptor agonist) showed a significative rescue of L3 larvae motility in TBPH hypomorphic background-null alleles compared to controls (Fig. 4C).

## CONCLUSIONS

Alterations in GABA neurotransmission were previously described in patients suffering from Alzheimer’s disease (Govindpani et al., 2017), Parkinson’s disease (Błaszczyk, 2016) and related neurodegenerative process including ALS (Foerster et al., 2013; Kim and Yoon, 2017). Nevertheless, the role of GABA in TDP-43 pathology was not clarified. In that respect we found that TBPH, the TDP-43 conserved protein in Drosophila, was required to maintain the correct intracellular levels of GABA. Regarding the molecular mechanisms involve, we discovered that TBPH regulates the expression levels of GAD1, the enzyme required to sustain GABA neurotransmitter expression, through the binding and processing of GAD1 mRNA (see also Romano et.al 2018). In addition, we observed that the pharmacological potentiation of GABA signaling with nipecotic acid, an inhibitor of GABA uptake, was able to significantly revert the locomotive defects observed in TBPH null flies. Altogether, the results reveal a novel molecular mechanism behind TDP-43-derived pathologies and put forward a potential pharmacological approach to compensate neurotransmitter balance defects in ALS affected individuals.

## DATA AVAILABILITY

The datasets generated during and/or analyzed during the current study are available from the corresponding author on reasonable request.

## ACKNOWLEDGEMENTS

We thank the Bloomington Stock Centre and Developmental Study Hybridoma Bank for stocks and reagents.

## Supplementary Material

**Supplementary Figure 1.**
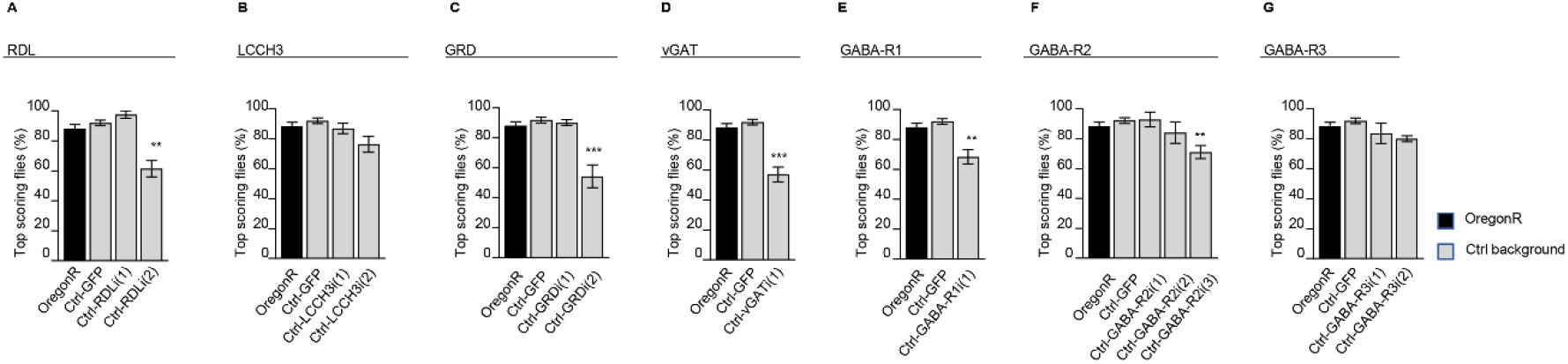
Silencing of GABA receptors in wild type background. **A-G** Climbing assay of 4 days adult flies of OregonR (black column), Ctrl-GFP (tbph^Δ23^,*elav*-GAL4/UAS-GFP;UAS-Dicer/+) and Ctrl-GABA receptor name (tbph^Δ23^,*elav*-GAL4/UAS-GABA-receptor IR;UAS-Dicer/+). In **A** RDL-IR line (1)= #41101, line (2) #100429; in **B** LCCH3-IR line (1) #37409, line (2) #109606; in **C** GRD-IR line (1) #38384, line (2) #58175; in **D** vGAT-IR line (1) #45916; in **E** GABA_B_ receptor type 1 line (1) #101440; in **F** GABA_B_ receptor type 2 line (1) #1784, line (2) #1785, line (3) #110268; in **G** GABA_B_ receptor type 3 line (1) #50622, line (2) #108036. The total number of tested animals per genotype was > 50. **p<0.01, ***p < 0.001; calculated by one-way ANOVA. Error bars SEM.

